# *C. elegans* heritably adapts to *P. vranovensis* infection via a mechanism that requires the cysteine synthases *cysl-1* and *cysl-2*

**DOI:** 10.1101/675132

**Authors:** Nicholas O. Burton, Cristian Riccio, Alexandra Dallaire, Jon Price, Benjamin Jenkins, Albert Koulman, Eric A. Miska

## Abstract

Parental exposure to pathogens can prime offspring immunity in diverse organisms. The mechanisms by which this heritable priming occurs are largely unknown. Here we report that the soil bacteria *Pseudomonas vranovensis* is a natural pathogen of the nematode *Caenorhabditis elegans* and that parental exposure of animals to *P. vranovensis* promotes offspring resistance to infection. Furthermore, we demonstrate a transgenerational enhancement of progeny survival when three consecutive generations of animals are exposed to *P. vranovensis.* By investigating the mechanisms by which animals heritably adapt to *P. vranovensis* infection, we found that parental infection by *P. vranovensis* results in increased expression of the cysteine synthases CYSL-1 and CYSL-2 and the regulator of hypoxia inducible factor RHY-1 in progeny and that these three genes are required for adaptation to *P. vranovensis.* To our knowledge, these observations represent the largest heritable increase in offspring survival in response to a pathogen infection reported in any organism to date and establish a new CYSL-1, CYSL-2, and RHY-1 dependent mechanism by which animals adapt to infection.

## Introduction

Intergenerational and transgenerational responses to environmental stress have been reported in evolutionarily diverse organisms^1–7^. These studies investigated a variety of types of stresses ranging from osmotic stress^1^, to mitochondrial stress^5^, to pathogen infection^8^. In each case, parental exposure to stress appeared to prime offspring to respond to a similar stress. For example, parental exposure of the nematode *Caenorhabditis elegans* to the opportunistic pathogen *Pseudomonas aeruginosa* was reported to alter offspring behavior in a way that promotes offspring avoidance of *P. aeruginosa* and enhances offspring survival^8^. Similarly, studies of *Arabadopsis thaliana* have demonstrated that parental exposure to mild osmotic stress can promote offspring resistance to future osmotic stress^9^. While many intergenerational and transgenerational effects of the environment have been described in plants and invertebrates, similar observations have recently been extended to vertebrates, including mammals. For example, parental exposure to high population density was demonstrated to promote an accelerated postnatal growth rate in red squirrels that enhanced offspring survival by allowing them to acquire territories more quickly^2^. Collectively, these findings raise the exciting possibility that parental exposure to environmental stress causing programmed changes in offspring physiology might represent a fundamental and significantly understudied aspect of inheritance with implications for diverse fields of biological and medical sciences.

Among the diverse environmental stresses an organism might encounter, pathogens such as viruses, bacteria, and some eukaryotes are among the most ubiquitous stresses found in nature^10^. To adapt to the constant threat of pathogens, many organisms have evolved mechanisms by which parental exposure to pathogens can prime offspring immunity^6, 11–16^. For example, in mammals, a mother can transfer specific antibodies to her offspring via milk to prime offspring immunity^16^. Similar observations of parents priming offspring immunity in response to pathogens have been reported in both plants^15^ and invertebrates^11^, even though these organisms lack antibodies. These findings suggest that multiple independent mechanisms have evolved for parents to prime offspring immunity.

The mechanisms by which parental exposure to pathogens might result in adaptive changes in offspring in organisms that lack antibodies remain largely unknown. Here we identify that *C. elegans* can heritably adapt to a natural pathogen, *Pseudomonas vranovensis*, and that adaptation to *P. vranovensis* requires the cysteine synthases *cysl-1* and *cysl-2*. Furthermore, we demonstrate that the exposure of animals to *P. vranovensis* can enhance the resistance of their progeny transgenerationally under conditions when three consecutive generations of animals are exposed to infection. We find that this heritable adaptation is not regulated by pathways previously reported to regulate transgenerational adaptations to environmental stress, suggesting that this heritable adaptation represents a new model of a transgenerational enhancement of survival in response to environmental stress^8^.

## Results

One of the major obstacles in determining the molecular mechanisms underlying the intergenerational and transgenerational effects of pathogen infection is the lack of robust models that can be easily analyzed in the laboratory. To address this obstacle, we sought to develop a robust model of the heritable effects of bacterial infection in the nematode *C. elegans* by testing whether parental infection of *C. elegans* with bacterial pathogens from its natural environment could affect offspring response to future infection. Previous sampling of bacterial species from the natural environments of *C. elegans* identified 49 as yet undescribed bacterial isolates that induce the expression of the immune response genes *irg-1* or *irg-5*^17^. We found that parental exposure to two of the isolates, BIGb446 and BIGb468, significantly enhanced offspring survival in response to future exposure to the same bacteria (Fig. 1A). Specifically, approximately 95% of newly hatched larvae from parents fed a standard laboratory diet of *E. coli* HB101 died within 24 hours of hatching on BIGb446 or BIGb468 (Fig. 1A). By contrast, 45% of embryos from adults briefly exposed to BIGb446 or BIGb468 were still alive at 24 hours (Fig. 1A) and a majority of these larvae survived through adulthood (Fig. 1B-C). In addition, we found that it took *P. vranovensis* approximately 72 hours to kill 95% of adult animals, indicating that adults are more resistant to BIGb446 than larval animals (Supplementary Fig. 1), and that UV-killed bacteria did not cause lethality, suggesting that live bacteria are required for killing (Fig. 1D). We conclude that parental exposure to bacterial isolates BIGb446 and BIGb468 enhances offspring survival in response to future exposure to these bacteria.

**Figure 1.**
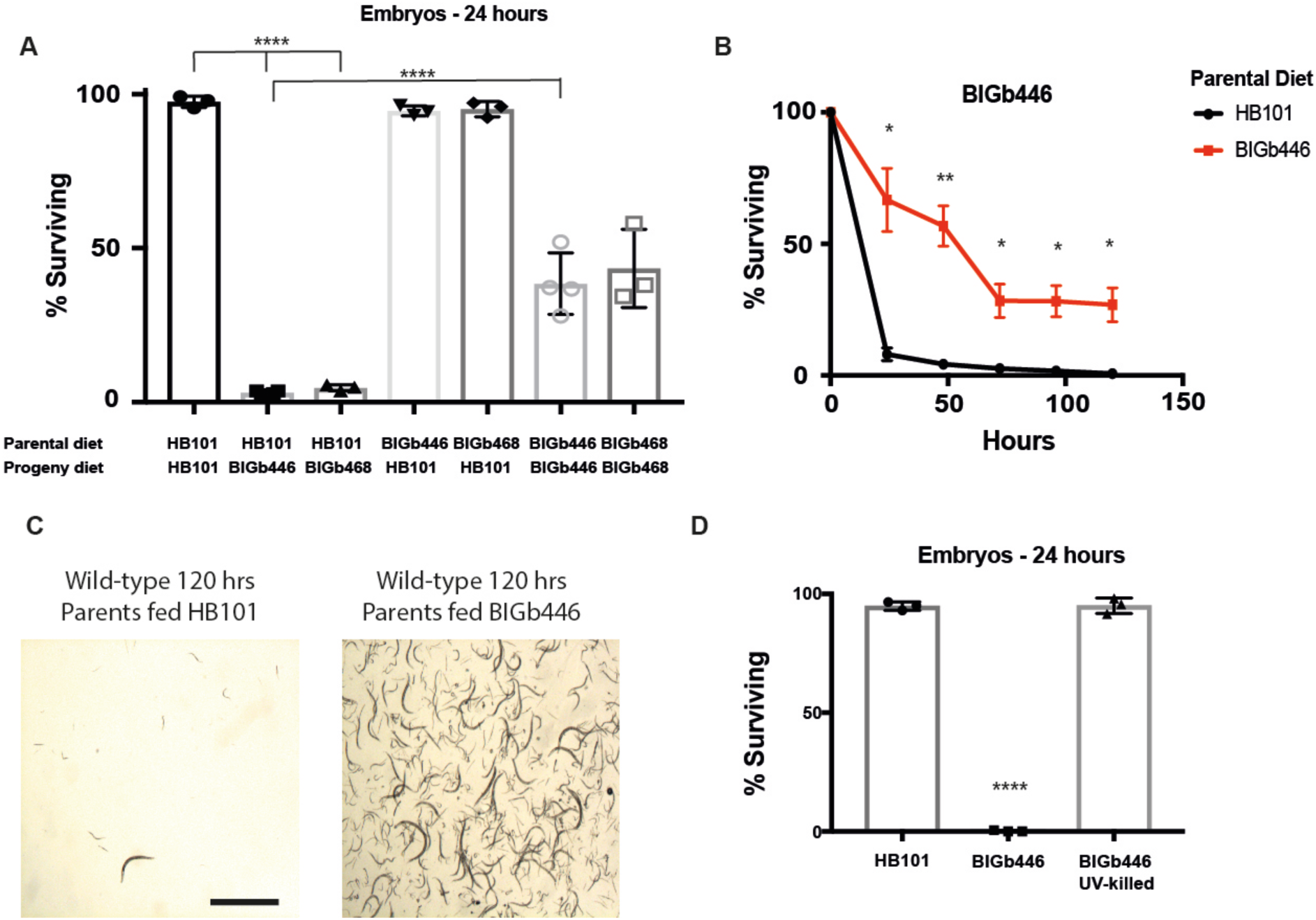
*C. elegans* heritably adapts to infection by *Pseudomonas vranovensis.* (A) Percent of wild-type animals surviving on plates seeded with either *E. coli* HB101 or bacterial isolates BIGb446 and BIGb468 after 24 hrs. Error bars, s.d. n = 3 experiments of >100 animals. (B) Percent of wild-type animals surviving on plates seeded with bacterial isolate BIGb446. Error bars, s.d. n = 3 experiments of 500 animals. (C) Images of wild-type animals surviving after 120 hrs of feeding on bacterial isolate BIGb446. 1000 animals were used at t = 0 in each condition and surviving animals were resuspended in 20 μl M9 and imaged. Scale bars 1 mm. (D) Percent of wild-type animals surviving on *E. coli* HB101 or bacterial isolate BIGb446 after 24 hrs. Error bars, s.d. n = 3 experiments of >100 animals. * = p < 0.05, ** = p < 0.01, *** = p < 0.001, **** p < 0.0001.

To determine the species identity of bacterial isolates BIGb446 and BIGb468 we performed long read whole genome sequencing and assembled the genomes of these bacteria. We found that the genomes of both BIGb446 and BIGb468 were approximately 5.9 Mb (Supplementary File 1 and 2) and were 99.38% identical across the entire genome (Supplementary Table 1 and Supplementary Fig. 2). We concluded that BIGb446 and BIGb468 are isolates of the same species. We compared the 16S rRNA sequence of BIGb446 to known bacterial genomes using BLAST. We found that the 16s rRNA sequence from BIGb446 is 99.93% identical to *Pseudomonas vranovensis* strain 15D11. Previous studies of *Pseudomonas* phylogeny have used the DNA sequences of *gyrB*, *rpoB*, and *recA* to differentiate species of *Pseudomonas,* with similarity above 97% set as the accepted species threshold^18, 19^. We found that the sequence of *gyrB* was 98.05% identical, the sequence of *rpoB* was 99.44% identical, and the sequence of *recA* was 98.66% identical to the sequences of homologous genes in *P. vranovensis* strain 15D11. No other species of *Pseudomonas* was greater than 97% identical to *Pseudomonas sp.* BIGb446 at any of these three genes. Furthermore, we compared the average nucleotide identity (ANI) of our assembly of *Pseudomonas sp.* BIGb446 with the genome of *P. vranovensis* strain 15D11 using OrthoANIu^20^. We found these two genomes had an average nucleotide identity of 97.33%. We conclude that *Pseudomonas sp.* BIGb446 and *Pseudomonas sp.* BIGb468 are isolates of *Pseudomonas vranovensis*, a Gram-negative soil bacteria^21^.

In some cases, the effects of parental stress on offspring have been reported to be intergenerational and only last a single generation^1^. In other cases, the effects of parental stress on offspring have been reported to persist transgenerationally and thus affect descendants several generations later^5, 8, 22^. To test whether exposure to *P. vranovensis* had any transgenerational effects on immunity we first fed adult animals *P. vranovensis* for 24 hours and assayed the response of progeny one and two generations later. We found that a single exposure of adult animals to *P. vranovensis* enhanced their offspring’s survival (Fig. 2A-B), an intergenerational effect (Fig. 2C), but did not enhance the survival of their descendants two generations later (Fig. 2A-B). These results indicate that a single generation of exposure to *P. vranovensis* only intergenerationally affect progeny survival (Fig. 2C).

**Figure 2.**
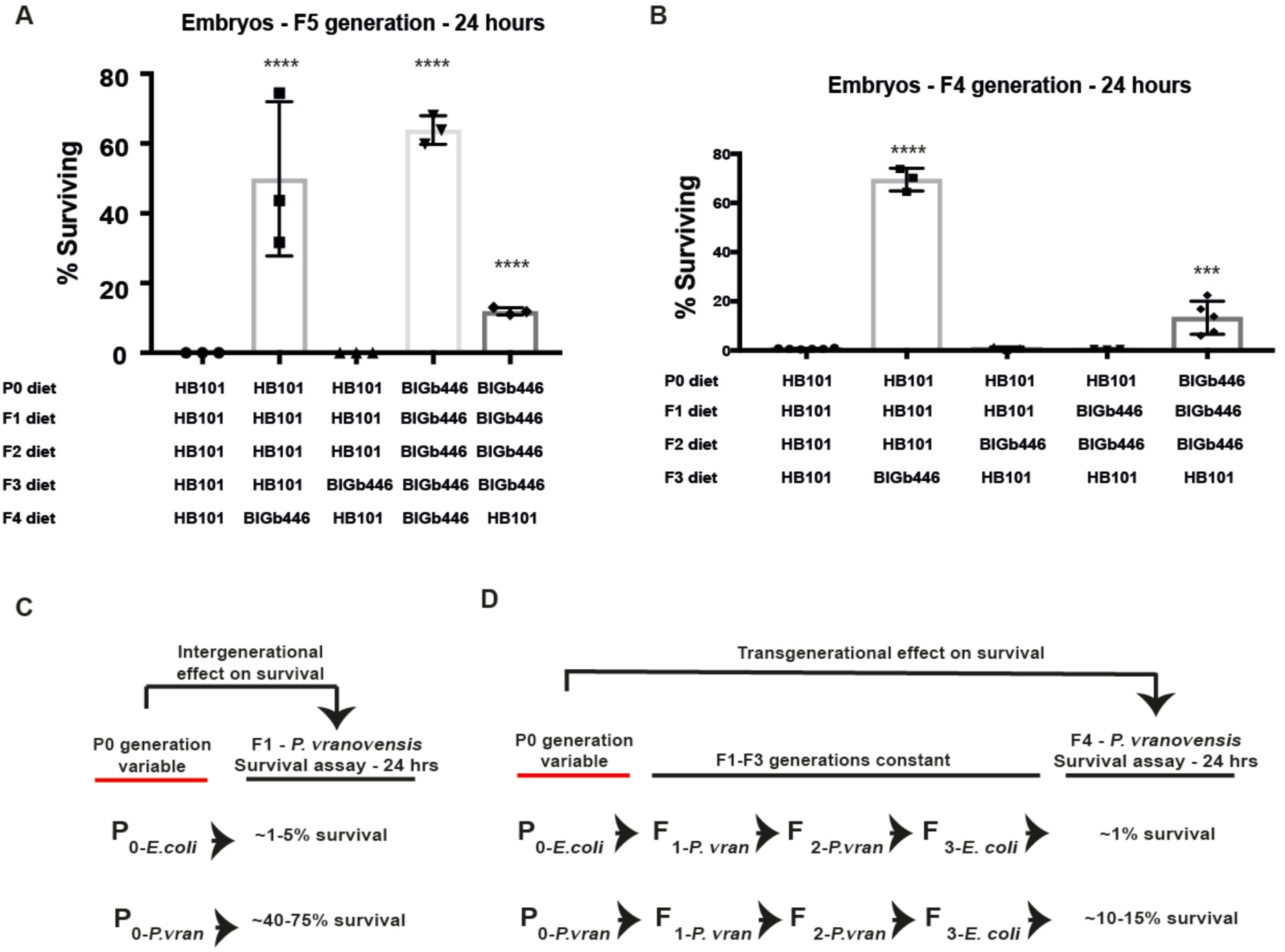
*C. elegans* adaptation to *P. vranovensis* can be inherited transgenerationally. (A) Percent of wild-type animals surviving on plates seeded with bacterial isolate BIGb446 after 24 hrs. Error bars s.d. n = 3 experiments of > 100 animals. (B) Percent of wild-type animals surviving on plates seeded with bacterial isolate BIGb446 after 24 hrs. Error bars s.d. n = 3 experiments of 500 animals. Wild-type animals carried the integrated transgene *nIs470.* (C) Diagram of intergenerational effects of *P. vranovensis* exposure on *C. elegans* survival. (D) Diagram of transgenerational effects of *P. vranovensis* exposure on *C. elegans* survival. Error bars s.d. n = 3 experiments of 500 animals. *** = p < 0.001, **** p < 0.0001.

Some studies of the effects of parental environment on offspring have found that multiple consecutive generations of exposure of animals to the same stress can enhance adaptation to stress in descendants^5^. We therefore tested whether the exposure of five consecutive generations (P0, F1, F2, F3, and F4) of animals to *P. vranovensis* could enhance F5 progeny survival in response to *P. vranovensis* infection but found that F5 survival in this case was not significantly different from F1 survival after a single generation of exposure to *P. vranovensis* (Fig. 2A). These results are consistent with *P. vranovensis* infection intergenerationally, but not transgenerationally, affecting progeny survival.

Finally, we tested whether the exposure of four consecutive generations of animals to *P. vranovensis* could cause animals’ resistance to *P. vranovensis* to persist for more than a single generation. Unlike a single generation of exposure to *P. vranovensis*, we found that the exposure of four consecutive generations of animals (P0, F1, F2, and F3) to *P. vranovensis* enhanced the survival of F5 animals, even when their F4 parents were not exposed to *P. vranovensis* (Fig. 2A). These data indicate multiple consecutive generations of exposure to *P. vranovensis* has different effects on the survival of descendants when compared to a single generation of exposure. Furthermore, these results suggest that multiple consecutive generations of exposure to *P. vranovensis* might have transgenerational effects on animal survival in response to future *P. vranovensis* infection.

Transgenerational effects are defined as an effect of the environment of P0 animals on F3 or later descendants^23^. To test whether multiple consecutive generations of exposure to *P. vranovensis* could have transgenerational effects on animal survival we exposed three (P0, F1, and F2), two (F1 and F2), and one (F2) generation of animals to *P. vranovensis* and assayed the survival of F4 generation animals exposed to *P. vranovensis*. We found that one generation of exposure (F2) and two consecutive generations of exposure to *P. vranovensis* (F1 and F2) did not affect F4 survival (Fig. 2B). By contrast, we found that three consecutive generations of exposure to *P. vranovensis* (P0, F1, and F2) did enhance the survival of F4 generation animals (Fig. 2B and 2D). These results indicate that the environment of P0 animals can enhance the survival of F4 generation animals under conditions where three or more generations of animals are exposed to *P. vraonvensis* (Fig. 2D). Furthermore, these results demonstrate that under conditions of multiple consecutive generations of animals being exposed to *P. vranovensis* the effects of exposure to *P. vranovensis* can transgenerationally affect progeny survival (Fig. 2D).

To better understand the mechanisms by which *C. elegans* can heritably adapt (defined by increased survival) to *P. vranovensis* we investigated how parental exposure to *P. vranovensis* can lead to a between 10 and 50-fold increase in offspring survival in response to future *P. vranovensis* exposure. In the case of this particular assay, the effect is intergenerational. However, we note that many previous studies of transgenerational effects in *C. elegans* have demonstrated that the same molecular mechanisms mediate both intergenerational and transgenerational effects of the environment, such as the inheritance of small RNA silencing^24^ and behavioral responses to *P. aeruginosa* infection^8^, suggesting that the molecular mechanisms that mediate many intergenerational and transgenerational effects of the environment might be the same. As a first approach, we tested whether mutations in any factors previously reported to be involved in intergenerational or transgenerational responses to stress were required for *C. elegans* heritable adaptation to *P. vranovensis* including small RNA mediated pathways (*hrde-1*^24^*, prg-1*^8^), H3K9 methylation (*set-32*^25^ and *met-2*^4^), H3K4 methylations (*set-2, wdr-5.1, spr-5*)^5, 22^, DNA adenosine methylation (*damt-1*)^5^, and the RAS/ERK signaling pathway (*lin-45*)^1^. We found that none of these factors were required for *C. elegans* adaptation to *P. vranovensis* (Fig. 3A) These data suggest that this adaptation to *P. vranovensis* involves an as-yet-unknown mechanism.

**Figure 3.**
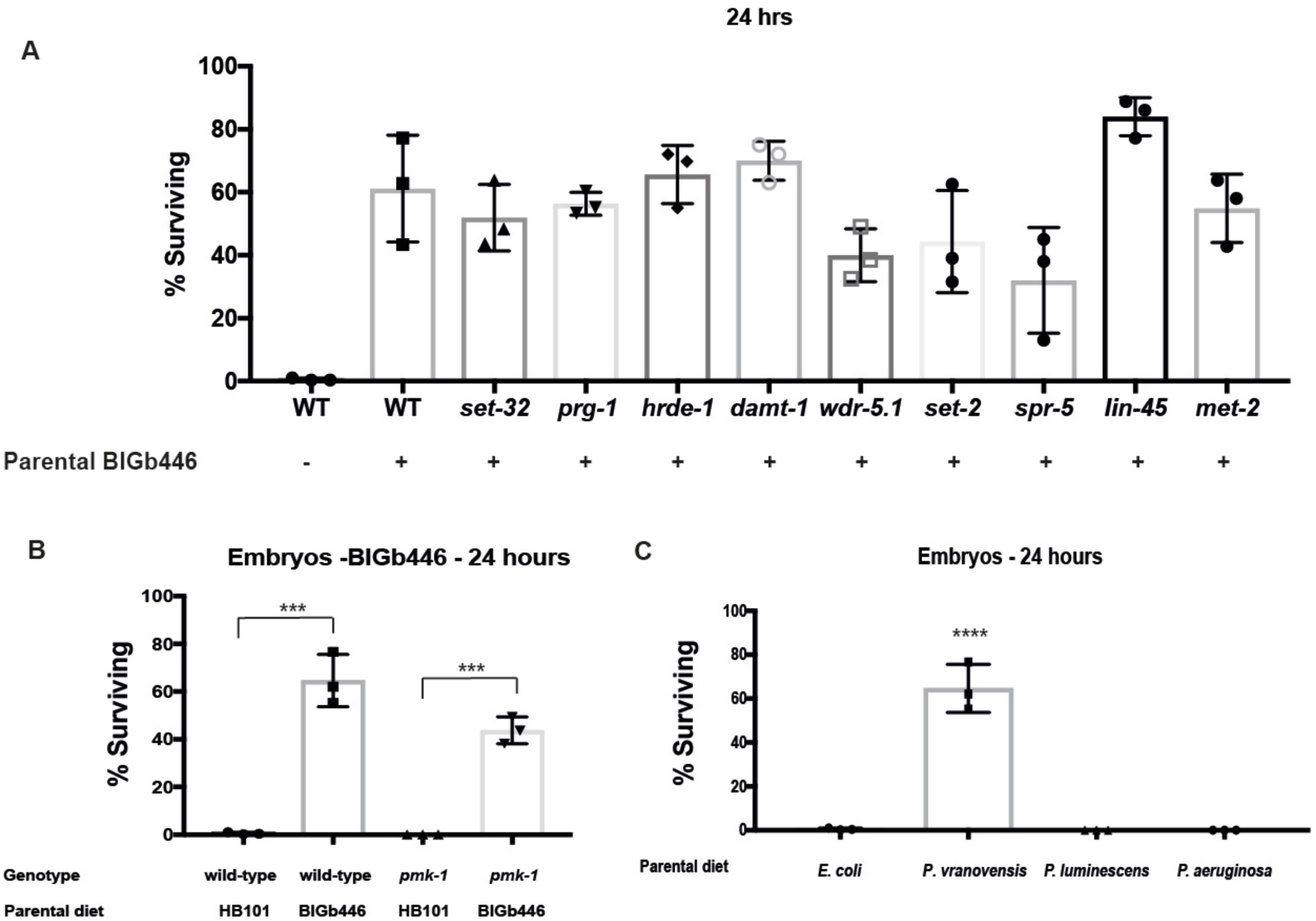
Adaptation to *P. vranovensis* does not require factors previously reported to be required for transgenerational effects in *C. elegans*. (A) Percent of wild-type, *set-32(ok1457), prg-1(n4357), hrde-1(tm1200), damt-1(gk961032), wdr-5.1(ok1417), set-2(ok952), spr-5(by134), lin-45(n2018),* and *met-2(n4256)* mutants surviving on plates seeded with bacterial isolates BIGb446 after 24 hrs. Error bars, s.d. n = 3 experiments of >100 animals. (B) Percent of wild-type and *pmk-1(km25)* mutants surviving on plates seeded with bacterial isolates BIGb446 after 24 hrs. Error bars, s.d. n = 3 experiments of >100 animals. (C) Percent of wild-type animals surviving on plates seeded with bacterial isolate BIGb446 after 24 hrs. Error bars s.d. n = 3 experiments of 500 animals. *** = p < 0.001, **** p < 0.0001.

Bacterial isolates BIGb446 and BIGb468 were originally reported to promote the expression of immune response gene *irg-5*^17^. The expression of *irg-5* in response to pathogen infection is controlled by the p38-like MAP kinase PMK-1^26^. We tested whether PMK-1 is required for resistance to *P. vranovensis* by placing wild-type and *pmk-1* mutant adults on plates seeded with *P. vranovensis*. We found that 100% of *pmk-1* mutants were dead after 48 hours while more than 40% of wild-type animals remained alive (Supplementary Fig. 1). These results indicate that PMK-1 promotes resistance to *P. vranovensis*. We tested whether PMK-1 was required for heritable adaptation to *P. vranovensis* by exposing wild-type and *pmk-1* mutants to *P. vranovensis* for 24 hours and assaying the survival of their offspring in response to repeated exposure to *P. vranovensis*. We found that PMK-1 is not required for the heritable adaptation to *P. vranovensis* (Fig. 3B).

*C. elegans’* heritable adaptation to *P. vranovensis* might be a general response to pathogen infection or specific to *P. vranovensis.* To test whether parental infection by other bacterial pathogens could protect offspring from *P. vranovensis* we exposed adult animals to two additional bacterial pathogens of *C. elegans*, *Pseudomonas aeruginosa* PA14^27^ and *Photorhabdus luminescens* Hb^28^, and assayed the response of their offspring to *P. vranovensis*. We found that parental infection by these two pathogens did not enhance offspring survival in response to *P. vranovensis* (Fig. 3C). These results indicate that adaptation to *P. vranovensis* is not a generic response to pathogenic bacteria.

Previous studies found that *C. elegans* can heritably adapt to osmotic stress and starvation by regulating gene expression and metabolism in offspring^1, 3^. To identify how parental infection by *P. vranovensis* affects offspring gene expression we profiled mRNA abundance by RNA-seq. We identified 1,153 genes that exhibit increased expression greater than 2-fold and 491 genes that decreased expression greater than 2-fold in embryos from parents exposed to *P. vranovensis* when compared to embryos from parents fed *E. coli* HB101 (Fig. 4A and Supplementary Table 2). We also quantified gene expression in young adults exposed to *P. vranovensis*. We found that 1,275 genes exhibited a greater than 2-fold change in expression when compared to young adults fed *E. coli* HB101 (Supplementary Table 2). Of these 1,275 genes, 398 were also observed to change in embryos (Fig. 4B). We conclude that infection by *P. vranovensis* results in similar, but distinct, effects on adult and embryonic gene expression.

**Figure 4.**
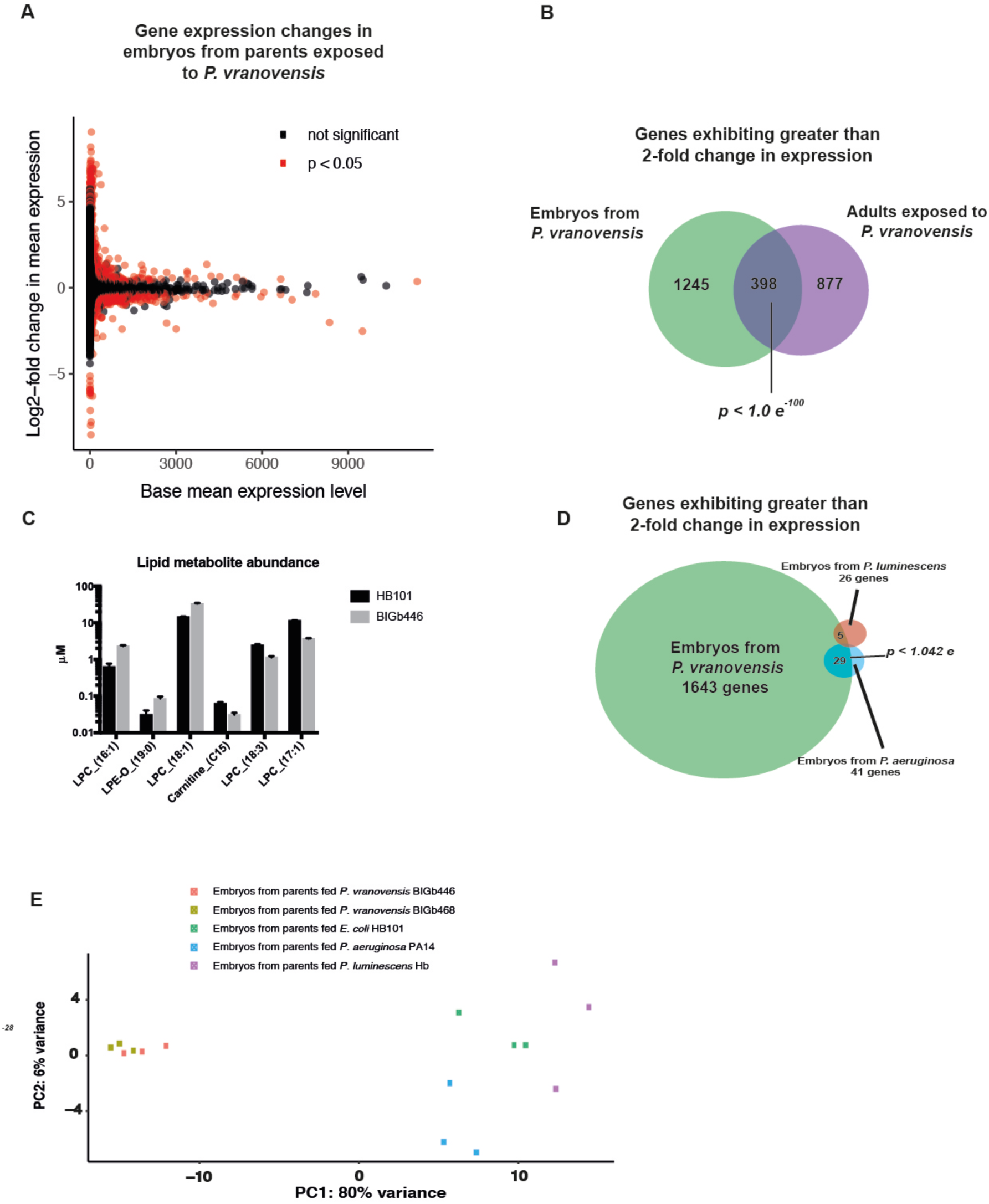
Parental infection by *P. vranovensis* alters gene expression and metabolism in offspring. (A) Gene expression changes in embryos from parents fed *P. vranovensis* BIGb446 when compared to embryos from parents fed *E. coli* HB101. Values represent averages from three replicates. (B) Venn diagram of genes exhibiting a greater than 2-fold change in RNA expression in adults fed *P. vranovensis* BIGb446 and embryos from adults fed *P. vranovensis* BIGb446. *p*-value represents normal approximation to the hypergeometric probability (See Statistics and Reproducibility) (C) μM abundance of lipid metabolites exhibiting a greater than 2-fold change in abundance in embryos from parents fed *P. vranovensis* BIGb446 when compared to embryos from parents fed *E. coli* HB101. Error bars, s.d. n = 3 replicates. (D) Venn diagram of genes exhibiting a greater than 2-fold change in RNA expression in embryos from parents fed *P. vranovensis* BIGb446, *P. aeruginosa* PA14, and *P. luminescens* Hb. *p*-value represents normal approximation to the hypergeometric probability (See Statistics and Reproducibility) (E) Principal component analysis (PCA) plot of mRNA expression data from RNA-seq of embryos from parents fed *E. coli* HB101, *P. vranovensis* BIGb446 and BIGb468, *P. aeruginosa* PA14, or *P. luminescens* Hb.

Previous studies of *C. elegans* found that parental exposure to osmotic stress can protect offspring from future exposure to osmotic stress by altering offspring metabolism^1^. We therefore tested whether parental exposure to *P. vranovensis* resulted in similar metabolic changes in offspring as parental exposure to osmotic stress. Using LC/MS we profiled 92 lipid metabolites, including those previously observed to heritably change in abundance in response to osmotic stress^29^. We found that only 6 metabolites exhibit modest changes in abundance in embryos from parents exposed to *P. vranovensis* when compared to embryos from parents fed *E. coli* HB101 (Fig. 4C and Supplementary Table 3). We conclude that parental exposure to *P. vranovensis* does not result in similar changes in offspring metabolism as parental exposure to osmotic stress^1^, and that these intergenerational adaptations to environmental stress are likely to be distinct.

We found that parental infection by *P. aeruginosa* and *P. luminescens* did not protect offspring from *P. vranovensis* (Fig. 3C). We hypothesized that different parental infections might result in different gene expression changes in offspring and that the specific changes caused by parental infection by *P. vranovensis* are required for offspring adaptation to *P. vranovensis*. To compare how parental infection by different bacterial pathogens affects offspring gene expression we exposed young adults to *P. aeruginosa* or *P. luminescens* for 24 hours and collected embryos from these animals to profile gene expression by RNA-seq. We found that only 41 genes exhibited a greater than 2-fold change in expression in embryos from parents exposed to *P. aeruginosa* when compared to embryos from parents fed *E. coli* HB101 and of these genes 29 also exhibited altered gene expression in embryos from parents exposed to *P. vranovensis* (Fig. 4D-E, Supplementary Table 4). Separately, we found that only 26 genes exhibited a greater than 2-fold change in expression in embryos from parents exposed to *P. luminescens* when compared to parents fed *E. coli* HB101 (Fig. 4D-E and Supplementary Table 4). Of these genes, only 5 also exhibited altered expression in embryos from parents exposed to *P. vranovensis* (Fig. 4D). Collectively, these results indicate that parental infection by *P. aeruginosa* and *P. luminescens* have only a limited effect on offspring gene expression when compared to parental infection by *P. vranovensis*. We conclude that parental exposure to different pathogenic diets has distinct effects on offspring gene expression.

A majority of genes that exhibit a greater than 2-fold change in expression in response to *P. vranovensis* exhibit an increase in mRNA expression (Supplementary Tables 2 and 5). Among the genes exhibiting the largest increase in expression in response to *P. vranovensis* we identified the cysteine synthases CYSL-1 and CYSL-2 (Fig. 5A-B and Supplementary Tables 2 and 5), which were previously reported to be involved in breaking down bacterial toxins produced by *P. aeruginosa*^30^. We confirmed that exposure of parents to *P. vranovensis* increases CYSL-2 expression in embryos using a GFP reporter (Fig. 5C)^31^. We hypothesized that increased expression of *cysl-1* and *cysl-2* in offspring might promote adaptation to *P. vranovensis*. We assayed seven independent alleles of *cysl-1* and two independent mutants lacking *cysl-2* and found that these mutants were unable to adapt to *P. vranovensis* (Fig. 5D-E and Supplementary Fig. 3). In addition, we found that the defect caused by loss of *cysl-2* could be rescued by expressing a wild-type copy of *cysl-2* (Fig. 5E). We conclude that CYSL-1 and CYSL-2 are required for adaptation to BIGb446.

**Figure 5.**
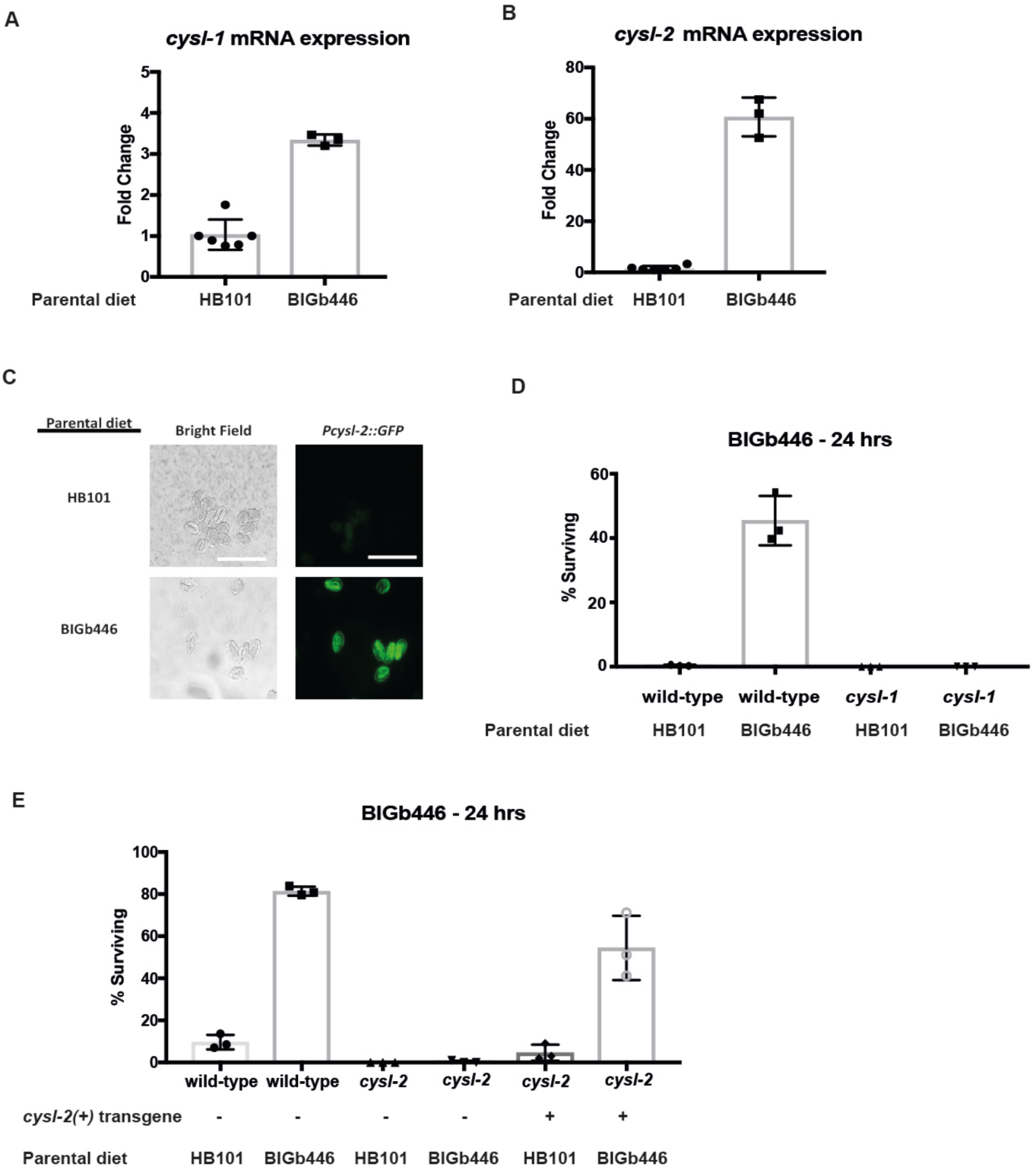
CYSL-1 and CYSL-2 are required for *C. elegans* to adapt to *P. vranovensis.* (A) Transcripts per million (TPM) of *cysl-1* in wild-type embryos from parents fed *E. coli* HB101 or *P. vraonvensis* BIGb446. Error bars, s.d. n = 3 replicates. (B) Transcripts per million (TPM) of *cysl-2* in wild-type embryos from parents fed *E. coli* HB101 or *P. vraonvensis* BIGb446. Error bars, s.d. n = 3 replicates. Data is the same as found in Table S2 and S5. (C) Representative images of *cysl-2::GFP* in embryos from parents fed *E. coli* HB101 or *P. vraonvensis* BIGb446. Scale bars 100 μm. (D) Percent of wild-type and *cysl-1(ok762)* mutants surviving on plates seeded with bacterial isolates BIGb446 after 24 hrs. Error bars, s.d. n = 3 experiments of >100 animals. (E) Percent of wild-type and *cysl-2(ok3516)* mutants surviving on plates seeded with bacterial isolates BIGb446 after 24 hrs. Error bars, s.d. n = 3 experiments of >100 animals. *** = p < 0.001, **** p < 0.0001.

CYSL-1 and CYSL-2 were previously found to function in a signaling pathway with the regulator of hypoxia factor, RHY-1, the EGLN1 homolog EGL-9, and the hypoxia inducible factor HIF-1 to regulate animals’ response to hypoxia ^31^. We found that one of these genes, *rhy-1*, was also among the genes that exhibited the largest increase in expression in response to *P. vranovensis* (Fig. 6A). We tested whether mutants lacking RHY-1, EGL-9, and HIF-1 also exhibited altered adaptation to *P. vranovensis*. We found that three independent mutations in *rhy-1* resulted in animals that did not adapt to *P. vranovensis* (Fig. 6B), similar to *cysl-1* and *cysl-2* mutants. This defect was rescued by expressing a wild-type copy of *rhy-1* (Fig. 6B). By contrast, we found that the loss of *hif-1* and *egl-9* did not affect adaptation to BIGb446 (Fig. 6C). These results indicate that RHY-1, CYSL-1, and CYSL-2 function separately from EGL-9 and HIF-1 to promote animals’ adaptation to *P. vranovensis*.

**Figure 6.**
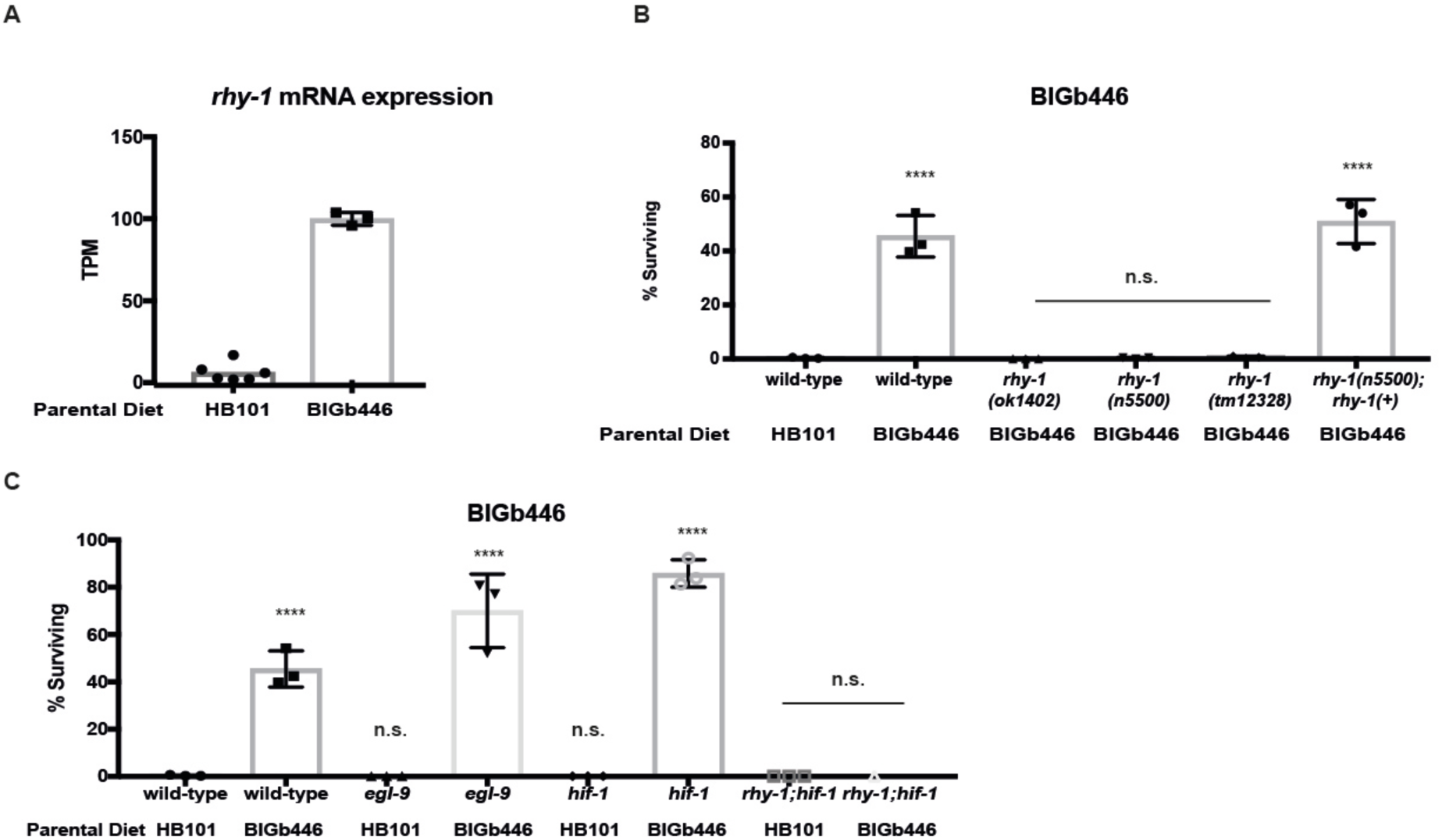
RHY-1 is required for *C. elegans* adaptation to *P. vranovensis*. (A) Transcripts per million (TPM) of *rhy-1* in wild-type embryos from parents fed *E. coli* HB101 or *P. vraonvensis* BIGb446. Error bars, s.d. n = 3 replicates. Data is the same as found in Table S2 and S5. (B) Percent of wild-type and *rhy-1* mutants surviving on plates seeded with bacterial isolates BIGb446 after 24 hrs. Error bars, s.d. n = 3 experiments of >100 animals. (C) Percent of wild-type, *egl-9(n586), hif-1(ia4),* and *rhy-1(n5500)* mutants surviving on plates seeded with bacterial isolates BIGb446 after 24 hrs. Error bars, s.d. n = 3 experiments of >100 animals. *** = p < 0.001, **** p < 0.0001.

## Discussion

Our results demonstrate that *P. vranovensis* is a pathogen of *C. elegans* and that parental exposure of *C. elegans* to *P. vranovensis* can protect offspring from future infection via a mechanism that requires the cysteine synthases *cysl-1* and *cysl-2* and the regulator of hypoxia inducible factor *rhy-1*. The ability of *P. vranovensis* to kill *C. elegans* larvae within 24 hours and the observation that *C. elegans* activates genes associated with oxidative stress in response to *P. vranovensis* suggests that *P. vranovensis* produces a toxic molecule(s) that is lethal to *C. elegans*. The cysteine synthases CYSL-1 and CYSL-2 have previously been reported to break down hydrogen cyanide^30^. These observations suggest that *C. elegans* might heritably adapt to infection *by P. vranovensis* by increasing the expression of CYSL-1 and CYSL-2 in offspring, which in turn protects offspring by breaking down hydrogen cyanide, a toxin that is known to be produced by other pathogenic species of *Pseudomonas*^32^. However, we note that *rhy-1* mutants were previously reported to be resistant to hydrogen cyanide mediated killing by *P. aeruginosa*^32^. By contrast, we found that *rhy-1* mutants are hypersensitive to *P. vranovensis* (Fig. 6B). These data suggest that resistance to potential hydrogen cyanide production by *P. vranovensis* alone does not explain the heritable adaptation to *P. vranovensis* and that our observations represent a new mechanism for *C. elegans* to adapt to a natural pathogen.

Recent studies of *C. elegans* response to the human opportunistic pathogen *P. aeruginosa* have also described a transgenerational response to this pathogenic species of *Pseudomonas*^8^. In this paradigm researchers found that exposure of a single generation of *C. elegans* to *P. aeruginosa* resulted in a heritable change in avoidance behaviour that persisted for four generations and was dependent on the PIWI Argonaute homolog PRG-1^8^. By contrast, we found that exposure of a single generation of *C. elegans* to *P. vranovensis* did not result in a transgenerational adaptation to infection, but rather the heritable increase in immunity only lasted one generation (Fig. 2A-B). Transgenerational effects of *P. vranovensis* infection were only observed after several consecutive generations of exposure to *P. vranovensis* (Fig. 2B and 2D). In addition, we found that *C. elegans* heritable adaptation to *P. vranovensis* did not require PRG-1 (Fig. 3A). The explanation for why *C. elegans* elicits different responses to *P. aeruginosa* when compared to *P. vranovensis* remains unclear. It is possible that *C. elegans* evolved distinct responses to *P. vranovensis* when compared to *P. aeruginosa*. Alternatively, studies of *C. elegans’* heritable responses to *P. aeruginosa* focused on changes in avoidance behaviour while our studies of *C. elegans* heritable responses to *P. vranovensis* focused on changes in survival under conditions where animals were unable to avoid the pathogen. It is possible that heritable changes in behaviour in response to bacterial infection in *C. elegans* are mediated by a small RNA pathway and PRG-1, while changes in immunity and the expression of cysteine synthases are controlled by a mechanism that is PRG-1 independent. Future studies will be important in differentiating between these possibilities and determining how parental infection can heritably prime offspring immunity, if these mechanisms are evolutionarily conserved, and how much they contribute to organisms’ responses to pathogens.

*C. elegans* was also recently reported to transgenerationally respond to infection by *P. aeruginosa* PAO1 and *Salmonella enterica* serovar Typhimurium strain MST1 by entering a stress resistant dauer developmental stage^33^. Entry into dauer only emerged after three consecutive generations (P0, F1, and F2) of exposure to these pathogens^33^. These results are similar to our observations of three consecutive generations of exposure to *P. vranovensis* could have transgenerational effects on progeny survival (Fig. 2). Taken together, these results suggest that some transgenerational effects of the environment, including adaptive effects, might only emerge after three consecutive generations of exposure to a particular stress.

Finally, studies of *C. elegans* interaction with the opportunistic pathogen *P. aeruginosa* indicate that parental exposure to *P. aeruginosa* can have both adaptive^8^ and deleterious^1^ consequences for offspring. We suspect that similar trade-offs, where adaptation to one stress comes at the cost of fitness in a different environment, are also likely to be the case for *C. elegans* adaptation to *P. vranovensis.* Future studies will likely be critical in determining what the costs of intergenerational and transgenerational responses to stress are, and we propose that such studies might be able to identify links between different types of stress responses. For example, previous studies have observed that *C. elegans* transcriptional response to osmotic stress and pathogen infection appear to be related^34^. These findings, in combination with observations that parental infection by *P. aeruginosa* results in offspring that are more susceptible to osmotic stress^1^, suggest that adapting to pathogen infection might generally disrupt animals’ ability to respond to osmotic stress and explain why animals are not normally resistant to this pathogen.

## Acknowledgments

We thank Buck Samuel, Gary Ruvkun, and Jonathan Ewbank for bacterial isolates; Long Ma, Keith Choe, Derek Sieburth, Bob Horvitz, Na An, and the *Caenorhabditis* Genetic Center, which is funded by the NIH National Center for Research Resources (NCRR), for strains; Martin Welch and George Salmond for use of bacterial culture equipment and space, and Martin Hemberg for feedback on RNA-seq analysis. N.O.B is funded by a Next Generation Fellowship from the Centre for Trophoblast Research. C.R. was supported by a Wellcome PhD programme. A.K. and B.J are funded by BBSRC grant BB/M027252/1. This work was also supported by Cancer Research UK (C13474/A18583, C6946/A14492) and the Wellcome Trust (104640/Z/14/Z, 092096/Z/10/Z) grants to E.A.M.

## Author Contributions

N.O.B. conceived the project and designed the experiments. N.O.B and E.A.M analysed the data. N.O.B., C. R., A.D., J.P., A.K., and B.J. performed the experiments. N.O.B. wrote the manuscript.

## Data and materials availability

RNA-seq data that support the findings of this study have been deposited at the European Nucleotide Archive (ENA) under the accession code PRJEB32993. The raw data for assembling the genomes of *P. vranovensis* isolate BIGb446 is available at the ENA under the accession code ERS3670403 and BIGb468 is available under the accession code ERS3670404. The raw data related to metabolomics is available on Dryad using doi:10.5061/dryad.8t6q5f5. Raw data supporting all figures is provided in Supplementary Table 6.

## Methods

### Strains

*C. elegans* strains were cultured as described^35^ and maintained at 20 °C unless noted otherwise. The Bristol strain N2 was the wild-type strain.

**LGI:** set-32(ok1457), prg-1(n4357), spr-5(by134)

**LGII:** cysl-2(ok3516, syb1431), damt-1(gk961032), rhy-1(n5500, ok1402, tm12398))

**LGIII:** wdr-5.1(ok1417), hrde-1(tm1200), set-2(ok952), met-2(n4256)

**LGIV:** pmk-1(km25), pgl-1(bn102), lin-45(n2018), nIs470[cysl-2::Venus; myo-2::RFP]

**LGV:** egl-9(n586), hif-1(ia4)

**LGX:** cysl-1(ok762, mr23, mr25, mr26, mr29, mr39, mr40)

**Unknown linkage:** burIs2[gst-31::gfp]

**Extrachromosomal arrays:** nEx1763[rhy-1(+); myo-2::RFP], burEx1 [cysl-2(+); myo-3::RFP]

### Sequencing of Pseudomonas vranovensis BIGb446 and BIGb468

Genomic DNA was prepped using a Gentra Puregene kit (QIAGEN). DNA was sheared to 10 kb using gTUBE (Covaris). Sheared DNA was barcoded and multiplexed for PacBio sequencing using Template Prep Kit 1.0-SPv3 and Barcoded Adapter Kit 8A (PacBio). Genomic DNA was sequenced using the PacBio Sequel system using version 3.0 sequencing reagents (PacBio) and 1M v3 SMRT cell.

### Genome assembly of BIGb446 and BIGb468

The genomes of BIGb446 and BIGb46 were assembled using HGAP4 from SMRT Link version 5.1.0.26412 with estimated genome sizes of 5 MB.

### Assays of adaptation to Pseudomonas vranovensis BIGb446 and BIGb468

*P. vranovensis* BIGb446 and BIGb468 was cultured in LB at 37 °C overnight. 1 mL of overnight culture was seeded onto 50 mm NGM agar plates and dried in a laminar flow hood (bacterial lawns completely covered the plate such that animals could not avoid the pathogen). All plates seeded with BIGb446 or BIGb468 were used the same day they were seeded. Young adult animals were placed onto 50 mm NGM agar plates seeded with 1 mL either *E. coli* HB101 or *P. vranovensis* BIGb446 or BIGb468 for 24 hours at room temperature. Embryos from these animals were collected and placed onto fresh NGM agar plates seeded with BIGb446 or BIGb468. Percent surviving were counted after 24 hours at room temperature unless otherwise noted.

### Transgenerational adaptation to P. vranovensis

*P. vranovensis* BIGb446 was cultured in LB at 37 °C overnight. 1 mL of overnight culture was seeded onto 50 mm NGM agar plates and dried in a laminar flow hood (bacterial lawns completely covered the plate). All plates seeded with BIGb446 were used the same day they were seeded. All animals in each generation were grown from embryos to young adults on NGM agar plates seeded with *E. coli* HB101. Young adults were then moved to fresh plates seeded with BIGb446 at room temperature for 24 hours. 24 hours of feeding on BIGb446 was counted as one generation of exposure to a BIGb446 diet. Embryos from parents fed BIGb446 were collected and placed onto plates seeded with *E. coli* HB101 until they were young adults and then moved to fresh plates seeded with BIGb446 for additional generations.

### Imaging of wild-type animals surviving after P. vranovensis exposure

1,000 embryos were placed onto NGM agar plates seeded with *P. vranovensis* BIGb446 at t = 0 in each condition and surviving animals at 120 hours were washed off of plates and resuspended in 20 μl M9 and imaged.

### Assays of C. elegans response to P. aeruginosa and P. luminescens

Young adult animals were placed onto slow-killing assay (*P. aeruginosa*) plates or NGM agar (*P. luminescens*) plates seeded with either *P. aeruginosa* PA14 or *P. luminescens* Hb for 24 hours at room temperature. Embryos from these animals were collected and snap frozen in liquid nitrogen for RNA sequencing and metabolomics analysis or placed onto fresh NGM agar plates seeded with BIGb446. Percent of animals placed on BIGb446 surviving were counted after 24 hours at room temperature.

### Assay of adult survival

Greater than 100 young adult animals were placed onto placed onto NGM agar plates seeded with *P. vranovensis* BIGb446 from overnight cultures in LB. Percent of animals surviving was counted at 24 hour intervals. Animals were scored as alive if they were mobile and dead if they were immobile and did not respond to touch.

### RNA-seq

Young adult animals were placed onto NGM agar plates seeded with *E. coli* HB101, *P. vranovensis* BIGb446 or BIGb468, or *P. luminescens* Hb or slow-killing assay plates seeded with *P. aeruginosa* PA14 for 24 hours at room temperature. Adult animals or embryos collected from adult animals after 24 hours were snap frozen in liquid nitrogen. Samples were lysed using a BeadBug microtube homogenizer (Sigma) and 0.5 mm Zirconium beads (Sigma). RNA was extracted using a RNeasy Plus Mini kit (Qiagen). mRNA was enriched using an NEBNext rRNA Depletion kit (NEB). Libraries for sequencing were prepped using an NEBNext Ultra II Library prep kit for Illumina (NEB) and loaded for paired-end sequencing using the Illumina HiSeq 1500.

### cysl-2::GFP *imaging*

To image adults and embryos, young adult animals expressing *nIs470* were placed onto NGM agar plates seeded with BIGb446 at room temperature for 24 hours. Embryos were collected and immediately imaged using a Zeiss AXIO imager A1 microscope and a Hamamatsu ORCA-ER camera.

### UV-killing bacteria

1 mL of overnight culture of BIGb446 was seeded onto 50 mm NGM agar plates and dried in the laminar flow hood. Seeded plates were then exposed to 20 μW/cm^2^ for 1 hour. Complete killing of bacteria was confirmed by testing inoculations of bacteria in LB overnight.

### cysl-2 cloning and rescue

Genomic *cysl-2* DNA was amplified from wild-type animals using the primers ACGATTGGGTTGGCTGTAAG and GGTCGTACGTGTTCGTTGTG. Extrachromosomal arrays were generated by injecting the corresponding PCR fragment and co-injection marker into the gonad of one-day old adults at the specified concentrations. *nobEx1* was generated by injecting *cysl-2* genomic DNA at 20 ng/μl and *myo-3::RFP* was injected at 10 ng/μl. Final injection DNA concentration was brought up to 150 ng/μl using DNA ladder (1kb HyperLadder – Bioline).

### Generation of cysl-2 CRISPR alleles

*syb1431* was generated by SunyBiotech. *syb1431* contains a 50 bp frameshift deletion in the first exon with the following flanking sequences ACCGGTGGTGAGCTCATCGGAAACACCCCA and GGTAGAGTACATGAACCCTGCCTGCTC.

### LC/MS lipid profiling

*C. elegans* was prepared for LC-MS lipidomics and acyl-carnitine analysis as previously described ^36^ with minor modifications. Briefly, ∼40 µL of concentrated embryos were re-suspended in 100 µL of water, then 0.4 mL of chloroform was added to each sample followed by 0.2 mL of methanol containing the stable isotope labelled acyl-carnitine internal standards (Butyryl-L-carnitine_-d7_ at 5 µM and Hexadecanoyl-L-carnitine_-d3_ at 5 µM). The samples were then homogenised by vortexing then transferred into a 2 mL Eppendorf screw-cap tube. The original container was washed out with 0.5 mL of chloroform: methanol (2: 1, respectively) and added to the appropriate 2 mL Eppendorf screw-cap tube. This was followed by the addition of 150 µL of the following stable isotope labelled internal standards (approximately 10 to 50 µM in methanol): Ceramide_C16_d31_, LPC_(C14:0_d42_), PC_(C16:0_d31_ / C18:1), PE_(C16:0_d31_ / C18:1), PG_(C16:0_d31_ / C18:1), PI_(C16:0_d31_ / C18:1), PS_(C16:0_d62_), SM_(C16:0_d31_), TG_(45:0_d29_) and TG_(48:0_d31_). Then, 400 µL of sterile water was added. The samples were vortexed for 1 min, and then centrifuged at ∼20,000 rpm for 5 minutes.

For the intact lipid sample preparation, 0.3 mL of the organic layer (the lower chloroform layer) was collected into a 2 mL amber glass vial (Agilent Technologies, Santa Clara California, USA) and dried down to dryness in an Eppendorf Concentrator Plus system (Eppendorf, Stevenage, UK) run for 60 minutes at 45 °C. The dried lipid samples were then reconstituted with 100 µL of 2:1:1 solution of propan-2-ol, acetonitrile and water, respectively, and then vortexed thoroughly. The lipid samples were then transferred into a 300 μL low-volume vial insert inside a 2 mL amber glass auto-sample vial ready for liquid chromatography separation with mass spectrometry detection (LC-MS) of intact lipid species.

For the acyl-carnitine sample preparation, 0.2 mL of the organic layer (the lower chloroform layer) and 0.2 mL of the aqueous layer (the top water layer) were mixed into a 2 mL amber glass vial and dried down to dryness. The dried acyl-carnitine samples were then reconstituted with 100 µL of water and acetonitrile (4: 1, respectively) and thoroughly vortexed. The acyl-carnitine samples were then transferred into a 300 μL low-volume vial insert inside a 2 mL amber glass auto-sample vial ready for liquid chromatography separation with mass spectrometry detection (LC-MS) of the acyl-carnitine species.

Full chromatographic separation of intact lipids ^37^ was achieved using Shimadzu HPLC System (Shimadzu UK Limited, Milton Keynes, United Kingdom) with the injection of 10 µL onto a Waters Acquity UPLC® CSH C18 column; 1.7 µm, I.D. 2.1 mm X 50 mm, maintained at 55 °C. Mobile phase A was 6:4, acetonitrile and water with 10 mM ammonium formate. Mobile phase B was 9:1, propan-2-ol and acetonitrile with 10 mM ammonium formate. The flow was maintained at 500 µL per minute through the following gradient: 0.00 minutes_40% mobile phase B; 0.40 minutes_43% mobile phase B; 0.45 minutes_50% mobile phase B; 2.40 minutes_54% mobile phase B; 2.45 minutes_70% mobile phase B; 7.00 minutes_99% mobile phase B; 8.00 minutes_99% mobile phase B; 8.3 minutes_40% mobile phase B; 10 minutes_40% mobile phase B; 10.00 minutes_40% mobile phase B. The sample injection needle was washed using 9:1, 2-propan-2-ol and acetonitrile with 0.1 % formic acid. The mass spectrometer used was the Thermo Scientific Exactive Orbitrap with a heated electrospray ionisation source (Thermo Fisher Scientific, Hemel Hempstead, UK). The mass spectrometer was calibrated immediately before sample analysis using positive and negative ionisation calibration solution (recommended by Thermo Scientific). Additionally, the heated electrospray ionisation source was optimised at 50:50 mobile phase A to mobile phase B for spray stability (capillary temperature; 380 °C, source heater temperature; 420 °C, sheath gas flow; 60 (arbitrary), auxiliary gas flow; 20 (arbitrary), sweep gas; 5 (arbitrary), source voltage; 3.5 kV. The mass spectrometer resolution was set to 25,000 with a full-scan range of m/z 100 to 1,800 Da, with continuous switching between positive and negative mode. Lipid quantification was achieved using the area under the curve (AUC) of the corresponding high resolution extracted ion chromatogram (with a window of ± 8 ppm) at the indicative retention time. The lipid analyte AUC relative to the associated internal standard AUC for that lipid class was used to semi-quantify and correct for any extraction/instrument variation.

Acyl-carnitine chromatographic separation was achieved using an ACE Excel 2 C18-PFP (150 mm, I.D. 2.1 mm, 2 µm) LC-column with a Shimadzu UPLC system. The column was maintained at 55 °C with a flow rate of 0.5 mL/min. A binary mobile phase system was used with mobile phase A; water (with 0.1% formic acid), and mobile phase B; acetonitrile (with 0.1% formic acid). The gradient profile was as follows; at 0 minutes_0% mobile phase B, at 0.5 minutes_100% mobile phase B, at 5.5 minutes_100% mobile phase B, at 5.51 minutes_0% mobiles phase B, at 7 minutes_0% mobile phase B. Mass spectrometry detection was performed on a Thermo Exactive orbitrap mass spectrometer operating in positive ion mode. Heated electrospray source was used; the sheath gas was set to 40 (arbitrary units), the aux gas set to 15 (arbitrary units) and the capillary temperature set to 250°C. The instrument was operated in full scan mode from m/z 75–1000 Da. Acyl-carnitine quantification was achieved using the area under the curve (AUC) of the corresponding high resolution extracted ion chromatogram (with a window of ± 8 ppm) at the indicative retention time. The acyl-carnitine analyte AUC relative to the associated internal standard AUC was used to semi-quantify and correct for any extraction/instrument variation. All lipid values were normalized to total lipid.

### RNA-seq and principal component analysis

Cutadapt version 1.18 was used to remove adapter sequences (AATGATACGGCGACCACCGAGATCTACACTCTTTCCCTACACGACGCTCTTCCG ATC for HS678 and AGATCGGAAGAGCACACGTCTGAACTCCAGTCA (forward) and AGATCGGAAGAGCGTCGTGTAGGGAAAGAGTGT (reverse) for HS755. Cutadapt was also used to trim the 3’ ends of reads when the phred quality value was below 20. Reads shorter than 40 bp were discarded. The genome sequence in FASTA format and annotation file in GFF3 format of *C. elegans* were downloaded from Ensembl release 96. The genome sequence was index with the annotation with hisat2 version 2.1.0. hisat2 was used for aligning reads to the reference genome and the maximum number of alignments to be reported per read was set to 5000. Featurecounts from the conda subread package version 1.6.3 was used to count the number of reads per gene. The Ensembl release 96 annotation file in GTF format for *C. elegans* was used. Fragments were counted when they overlapped an exon by at least 1 nucleotide and fragments are reported at the gene level. The option for the stranded protocol was turned off. Only read pairs that had both ends aligned were counted. Given our average fragment length of 300 bp, a distance of between 50 and 600 nucleotides was tolerated for read pairs. Read pairs that had their two ends mapping to different chromosomes or mapping to the same chromosome but on different strands were not counted. Multi-mapping reads were not counted. The raw counts table was imported into R 3.5.1 for differential expression analysis with DESeq2 version 1.22.1 Normalisation was carried out within each contrast. The PCA plots were produced with DESeq2 function plotPCA() after variance stabilising transformation of the data. A snakemake workflow (CITE snakemake ^38^) was created for the RNA-seq analysis and can it be found at https://github.com/cristianriccio/celegans-pathogen-adaptation.

### Genomic DNA alignment

MUMmer (*Version 3.0, default parameters*) and MUMmer-plot^39^ were used to visualize global alignments of BIGb446 and BIGb468 whole genome assemblies. Further to this NUCmer ^39^ and the associated dnadiff script was used to produce statistics on the alignment of the two bacterial genomes.

### Statistics and reproducibility

ANOVA analysis with post hoc p-value calculations was used for Fig. 1a, 1d, 2a, 2b, 2c, 2d, 2e, 4d, 5c, 5d, 6b, 6c, and Supplementary Fig. 3. Two-tail t-tests were used for Fig. 1b and Supplementary Fig. 1a. For Fig. 3b, 3d, and 4e the *p*-value was calculated using a normal approximation to the hypergeometric probability, as in http://nemates.org/MA/progs/representation.stats.html. For lipidomics data (Fig. 3c and Supplementary Table 3) p-values were calculated using two-tail t-tests and statistical significance was determined using Bonferonni correction for multiple hypotheses. * = p < 0.05, ** = p < 0.01, *** = p < 0.001, **** p < 0.0001. No statistical method was used to predetermine sample size. Sample sizes were chosen based on similar studies in the relevant literature. The experiments were not randomized. The investigators were not blinded to allocation during experiments and outcome assessment.

---

**Supplementary Figure 1.**
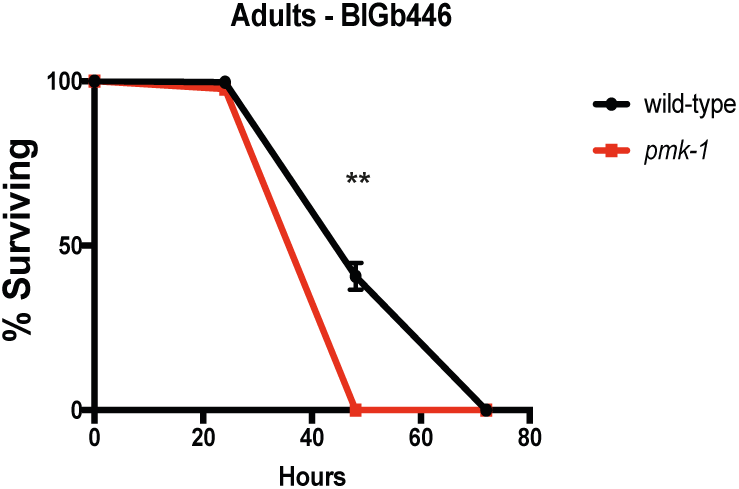
Genes associated with transgenerational inheritance and pathogen infection in *C. elegans* are not required for heritable adaptation to *P. vranovensis* BIGb446. Percent of wild-type and *pmk-1(km25)* mutant adults surviving on NGM plates seeded with P. vranovenis BIGb446. Error bars, s.d. n = 3 replicates of 100 animals. ** = p < 0.01

**Supplementary Figure 2.**
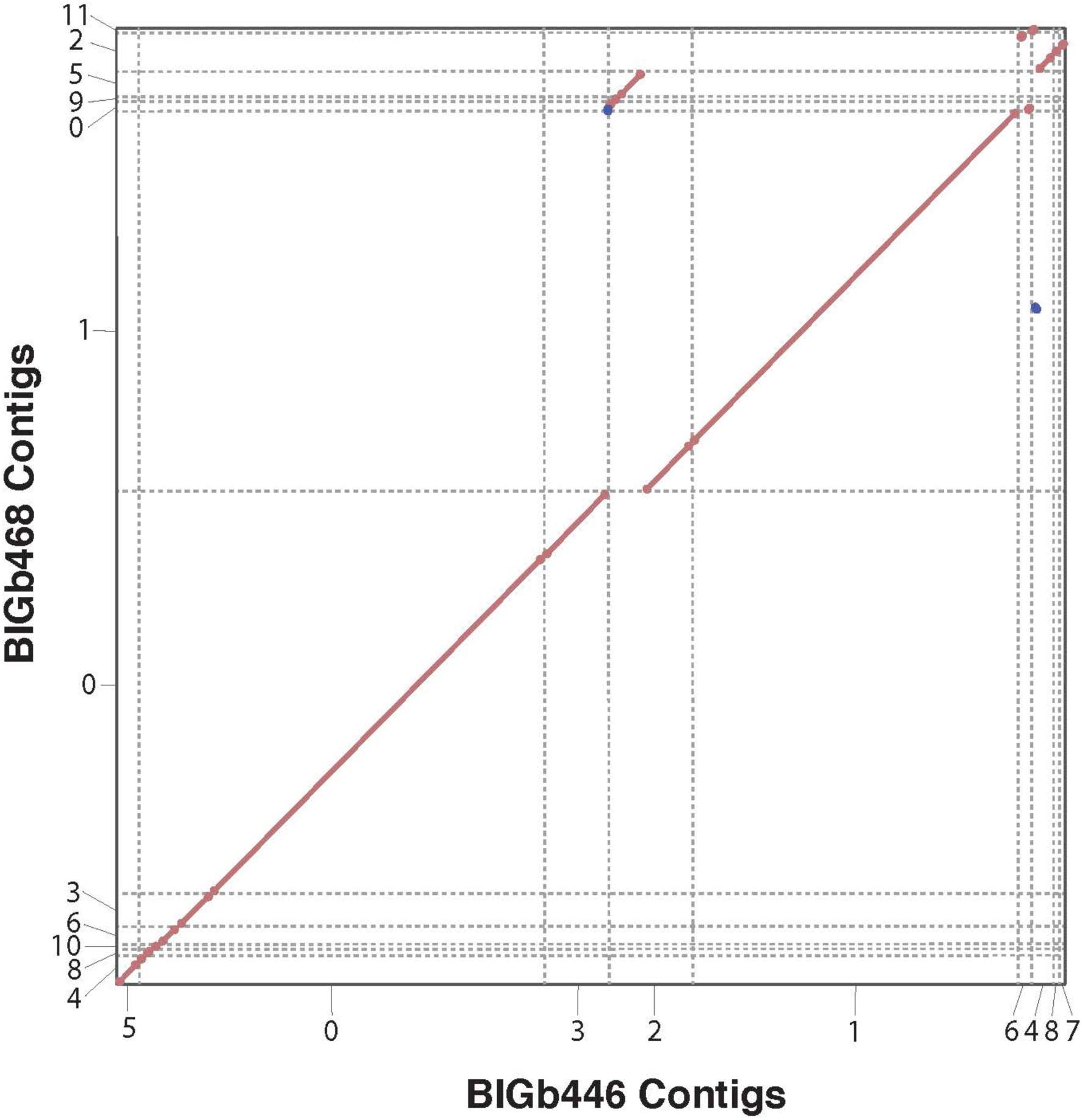
BIGb446 and BIGb468 are isolates of a single species. MUMmer plot of the alignments of contigs of BIGb446 and BIGb468 assembled genomes.

**Supplementary Figure 3.**
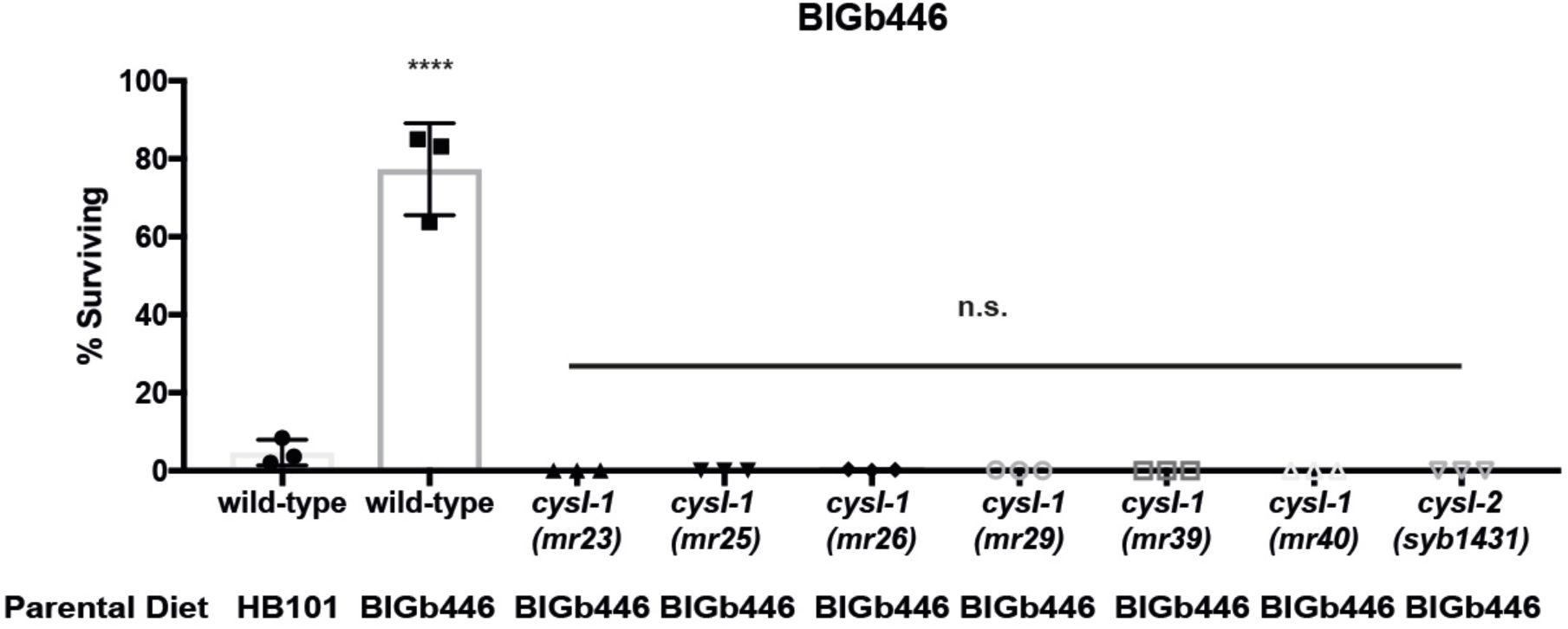
CYSL-1 and CYSL-2 are required for adaptation to *P. vranovensis* BIGb446. Percent of wild-type and *cysl-1(mr23, mr25, mr26, mr29, mr39, mr40)* and *cysl-2(syb1431)* mutants surviving on plates seeded with bacterial isolates BIGb446 after 24 hrs. Error bars, s.d. n = 3 experiments of >100 animals. **** p < 0.0001.

Supplementary File 1. Genome assembly of *P. vranovensis* BIGb446. FASTA format file of *P. vranovensis* BIGb446 genomic DNA.

Supplementary File 2. Genome assembly of *P. vranovensis* BIGb468. FASTA format file of *P. vranovensis* BIGb468 genomic DNA.

Supplementary Table 1. Comparison of genomic DNA sequences of BIGb446 and BIGb468 to *P. vranovensis*.

Supplementary Table 2. DEseq2 and TPM analysis of mRNA expression in wild-type embryos from parents fed *E. coli* HB101 or *P. vranovensis* BIGb446.

Supplementary Table 3. Profile of lipid metabolite abundances in wild-type embryos from parents fed *E. coli* HB101 or *P. vranovensis* BIGb446.

Supplementary Table 4. DEseq2 analysis of mRNA expression in wild-type embryos from parents fed *E. coli* HB101, *P. aeruginosa* PA14, or *P. luminescens* Hb.

Supplementary Table 5. Statistics Source Data

